# Bimodal electroencephalography-functional magnetic resonance imaging dataset for inner-speech recognition

**DOI:** 10.1101/2022.05.24.492109

**Authors:** Foteini Simistira Liwicki, Vibha Gupta, Rajkumar Saini, Kanjar De, Nosheen Abid, Sumit Rakesh, Scott Wellington, Holly Wilson, Marcus Liwicki, Johan Eriksson

## Abstract

The recognition of inner speech, which could give a ‘voice’ to patients that have no ability to speak or move, is a challenge for brain-computer interfaces (BCIs). A shortcoming of the available datasets is that they do not combine modalities to increase the performance of inner speech recognition. Multimodal datasets of brain data enable the fusion of neuroimaging modalities with complimentary properties, such as the high spatial resolution of functional magnetic resonance imaging (fMRI) and the temporal resolution of electroencephalography (EEG), and therefore are promising for decoding inner speech. This paper presents the first publicly available bimodal dataset containing EEG and fMRI data acquired nonsimultaneously during inner-speech production. Data were obtained from four healthy, right-handed participants during an inner-speech task with words in either a social or numerical category. Each of the 8-word stimuli were assessed with 40 trials, resulting in 320 trials in each modality for each participant. The aim of this work is to provide a publicly available bimodal dataset on inner speech, contributing towards speech prostheses.

## Background & Summary

Although research in the field of brain-computer interfaces (BCIs) began in the 1960s, it has accelerated in recent years due to advances in machine learning, imaging, and other data collection modalities.^1, 2^. A core aim of BCI research is to assist people who have lost the ability to move, speak or communicate with their environment. Inner speech can be described as the inner voice inside our heads; this phenomenon is used when thinking in a language without any accompanying muscle movement or speech articulation^3–6^. Decoding inner speech from brain activity is a burgeoning research area and has applications for BCI paradigms such as speech prostheses^7, 8^, in clinical contexts—for example, informing models of psychiatric disorders in which inner speech is disturbed (e.g., schizophrenia^9, 10^)—and in neuroscience, by deepening our understanding of the spatiotemporal neural dynamics of inner speech^11^.

Preliminary results have revealed that the most important parts of the brain for inner speech are the frontal gyri, including Broca’s area, the supplementary motor area and the precentral gyrus^12, 13^. Furthermore, core representations of the language system (phonology, lexicon, and syntax) have a clearly distinguishable spatial distribution in the neocortex^14–16^. This distribution of brain regions is remarkably similar across languages and across individuals^17^, regardless of why these language representations are accessed (i.e., for production or comprehension) or how they are accessed (i.e., visually (by reading) or auditorily (by listening)).

BCI technologies use brain data acquired by invasive (e.g., electrocorticography (ECoG)^18^) or noninvasive modalities (e.g., electroencephalography (EEG)^19^, functional magnetic resonance imaging (fMRI)^20^, functional near-infrared spectroscopy (fNIRS)^21, 22^ and magnetoencephalography (MEG)^23, 24^) to establish an interface between humans and machines; in particular EEG data are the most commonly used in BCIs; and fMRI is a typical complimentary modality due to high spatial resolution. BCI paradigms include motor imagery^25, 26^ and external stimulation paradigms, such as the visual P300^27^. In motor imagery paradigms, patients imagine their movement without overtly performing the action; in the visual P300 paradigm, patients typically use the direction of their eye gaze to spell out words by selecting among flashing stimuli, again without additional overt movement, which requires substantial participant concentration^28^. Thus, in recent years, the research focus for BCIs used to enable Augmentative and Alternative Communications (AACs) has turned to inner speech.

Research on inner speech decoding has investigated the use of all invasive ECoG^29, 30^ and noninvasive methods^31–35^. Various datasets have been acquired. Selected studies presenting EEG and fMRI are as follows: KARA ONE^36^ is a dataset of inner and outer speech recordings that combines a 62-channel EEG with facial and audio data. The dataset includes 12 participants, and the lexicon contains 7 phonemes and 4 phonetically-similar words for binary phonological classification. Coretto *et al*. ^37^ provided a dataset containing a 6-channel EEG recordings of inner and outer speech recordings of 5 vowels and 6 words. Nguyen *et al*. ^38^ generated a dataset that contains a 64-channel EEG recordings of inner speech from 15 subjects, with a lexicon of 3 vowels and 5 words (note that the work also introduces new algorithms, but this is secondary for our study). Ferreira *et al*. ^39^ provided a fMRI dataset of inner speech recordings from 20 native Portuguese speakers that consisted of cardinal vowels, monosyllabic and disyllabic words, and sentences. Recently, Nieto *et al*. ^40^ published an open-source unimodal EEG dataset of inner-speech BCI commands in Spanish.

The main limitation of such unimodal datasets is a much lower bound for possible recognition performance, as either temporal or spatial aspects of the data are not included. Unimodal datasets based on either EEG or fMRI can have drawbacks with regard to their temporal and spatial resolutions; specifically EEG datasets suffer from low spatial resolution but have a high temporal resolution, whereas fMRI datasets have a high spatial resolution that provides a deeper look into the subcortical structures of the brain but is limited by low temporal resolution. The relative strengths and weaknesses of these two neuroimaging modalities make their combination complimentary for brain analyses.

As for the combination of different modalities, recent studies on tasks different to inner-speech decoding have shown a possible improvement of the neural decoding performance^41–44^. Perronnet *et al*. ^41^ found that haemodynamic and electrophysiological activity during motor imagery tasks was higher when combining EEG and fMRI data compared to when EEG or fMRI data were used alone. Lioi *et al*.’s^43^ neurofeedback-based dataset of bimodal motor imagery was acquired with simultaneous EEG-fMRI recordings; the data were recorded from 30 subjects performing kinaesthetic motor-imagery tasks with the right hand to bring a ball to a target. In this work, the simultaneous bimodal EEG and fMRI dataset shows the potential of improving the quality of neurofeedback during a motor-imagery task compared to when using only one modality. Berezutskaya *et al*. ^44^ created a publicly-available multimodal nonsimultaneous dataset consisting of ECoG and fMRI data involving naturalistic simulation with a short audio-visual film; the dataset contains ECoG data from 51 subjects (5-55 years of age) and fMRI data from 30 participants (7-47 years of age) on the same task, enabling between-modality and subject-similarity analyses. This bimodal dataset shows the potential of combining different modalities to improve the study of neural mechanisms during language understanding and perception. The major outcomes of these studies were an improvement of the analysis when data from different modalities were combined. Making use of modality combination, hence, would be promising for inner-speech decoding as well.

The closest related work to this study is Cooney *et al*. ^42^, which generated a bimodal dataset of EEG (64-channel) and fNIRS (8-channel) data by acquiring simultaneous recordings from 19 subjects during outer and inner speech. However, the improvement in the performance in the task of inner-speech decoding was not as significant. Specifically, fNIRS showed a low decoding performance, and the use of the fNIRS modality was proven not significant for the bimodal decoding. Therefore, the choice of this work is to focus on EEG without fNIRS.

In terms of simultaneous vs nonsimultaneous recordings, we decided to choose nonsimultaneous recordings. Following the assessment procedure proposed by Scrivener^45^, we weighed in the following reasons: the analysis does not require simultaneously recorded data and it is not acceptable that the EEG data contain more artifacts when recorded with fMRI. Furthermore, nonsimultaneous recordings enable optimization of the task for each modality, such as fast paradigms with EEG and slow paradigms with fMRI, which has a slow BOLD response, therefore optimal for the bimodal acquisition of EEG and fMRI when comes to the inner-speech task.

The aim of this study was to collect separately-recorded EEG and fMRI recordings from healthy participants, performing an inner-speech task that followed the same experimental protocol for both modalities. This study showed that combining separately-recorded EEG and fMRI data can facilitate the decoding of inner speech, as this approach combines both high temporal and spatial resolution. To the best of our knowledge, this study represents the first publicly available dataset with bimodal nonsimultaneous EEG and fMRI recordings of inner speech. This bimodal dataset will allow future users to investigate the potential advantages of using bimodal *versus* unimodal data for inner-speech recognition and will also contribute towards the BCI development in the area of speech prostheses.

## Methods

### Participants

In order to identify participants for our study, we announced the study by distributing flyers describing the experimental procedure and aim of the study, at the Lulea University of Technology, following a list of predefined inclusion and exclusion criteria. In particular, the following inclusion criteria were followed: The current study aimed for an even gender distribution. In order to homogenize the sample, only right-handed people were considered in the study. To facilitate communication during data collection, which is mainly carried out by a non-Swedish-speaking person, primarily English speakers were consulted. In the same manner, the following exclusion criteria were followed: If people have difficulty understanding or following the instructions given at the time of preparation or if for some reason they were feeling uncomfortable during the magnetic resonance imaging (MRI) or EEG examination, they were excluded from the study. All participants filled out an fMRI pre-screening form in order to exclude people that should not undergo the experimental procedure (e.g., due to the presence of metallic objects in their body, claustrophobia etc.). The study was approved by the *Swedish Ethical Review Authority (Etikprövning myndigheden, ID:2021-06710-01*) (https://etikprovningsmyndigheten.se/) in accordance with the Swedish Ethical Review Act (SFS 2003:460). Ten participants filled out the questionnaire and as our ethical approval allowed only for a limited amount of subjects, we decided to include only right-handed subjects covering both genders (at least 40% of each gender). As a result, five healthy right-handed subjects aged 33-51 years, participated in this study (three females and two males). Detailed information on the subjects in this study is shown in Table 1. None of these subjects were native English speakers.

**Table 1.** Participant characteristics

| Participant | Self-declared sex | Age | Handedness | Native language |
| --- | --- | --- | --- | --- |
| sub-01 | Male | 33 | Right | Hindi |
| sub-02 | Male | 35 | Right | Bengali |
| sub-03 | Female | 51 | Right | Greek |
| sub-04 | Female | 35 | Right | Arabic |
| sub-05 | Female | 37 | Right | Hindi |

All subjects followed the same experimental protocol for the two modalities. The acquisition of the EEG and fMRI recording were performed sequentially followed the general approach of having an EEG recording followed by a fMRI recording with at least one hour of relaxation in between. In this study, we refer to the subjects with the following naming convention: sub-01, sub-02, sub-03, sub-04 and sub-05. Due to high fluctuations during the EEG recording the data from sub-04 were excluded from this study. All subjects provided written consent to participate in the study and to publish this dataset.

### fMRI hardware and setup

The data were collected using a Siemens Magnetom Prisma MRI system (Siemens Healthineers, Erlangen, Germany), equipped with a 20-channel head coil. The visual stimuli were presented during the fMRI recording from a computer to an Ultra HD LCD display (NordicNeuroLab, Bergen, Norway). The screen was 88×48 cm (3,840×2,160 pixels at full resolution).

Anatomical images were acquired using a sagittal T1-weighted 3D magnetization-prepared rapid acquisition gradient echo (MPRAGE) sequence with the following parameters: repetition time (TR) = 2.3 ms; echo time (TE) = 2.98 ms; inversion time (TI) = 900 ms; flip angle = 9°; slices = 208; matrix size = 256×256; and voxel size = 1×1×1 mm. Right after the anatomical scan, two field maps were obtained (A and B) with the following parameters: TR=662.0 ms, TE: A=4-92 ms, B=7.38 ms; and voxel size = 3×3×2 mm. Next, functional maps were obtained using double-echo gradient echo imaging BOLD sequences parallel to the bicommissural plane with the following parameters: TR = 2.16 s; TE = 30 ms; slices = 68; matrix size = 100×100 and voxel size = 2×2×2 mm.

### EEG hardware and setup

The EEG data were acquired using the *BioSemi Active2* measuring system (BioSemi B.V., Amsterdam, Netherlands) with a 16-bit resolution and a sampling rate of 512 Hz. A BioSemi EEG head cap with 64 electrodes in pre-fixed electrode positions and 6 external sensors was used. An appropriate cap size was selected for each participant by measuring his or her head circumference from nasion to inion. We also ensured that the cap was properly centred with the *Cz* (Vertex) at the centre of the head, namely, halfway between the nasion and inion and halfway between the two ears. *SignaGel* (Parker Laboratories BV, Almelo, Netherlands) was applied to each electrode to provide electrode connectivity with the subject’s head. All six external electrodes (EXG1-EXG6) were placed using stickers. The locations of the six electrodes were as follows:

- EXG1: On the left mastoid behind the left ear
- EXG2: On the right mastoid behind the right ear
- EXG3: 1 cm to the left of the left eye (aligned to the centre of the eye)
- EXG4: 2 cm above the left eye (aligned to the centre of the eye)
- EXG5: 2 cm below the right eye (aligned to the centre of the eye)
- EXG6: 1 cm to the right of the right eye (aligned to the centre of the eye)

EEG data were recorded with ActiView software, which was also developed by BioSemi. ActiView enables verification of the electrode impedance as well as the overall quality of the incoming data. The impedance of each electrode was manually examined at the beginning of each recording session, to ensure that it was between −20 μV and 20 μV; any electrodes not within this range were adjusted before recording to ensure the correct impedance, by adding/removing some gel, moving the participant’s hair underneath the electrode or wiggling the electrode. Lights in the room were dimmed to avoid subject’s eye flickering due to the high contrast between the room and the visual display.

### Experimental protocol

The overall experimental protocol consisted of two fMRI sessions and one EEG session and was performed over a period of 3 consecutive days. In this study, all EEG recordings were performed first, as the EEG setup can sometimes induce difficulties (e.g., achieving good electrode connectivity with the participant’s head). During day-01, the EEG recordings of sub-01, sub-02 and sub-03 took place followed by the recordings of the first fMRI session. The majority of the fMRI recordings for the second session were performed during day-02; only the recordings of sub-05 were performed on day-03. There was always a relaxation period of at least one hour in between the recordings and proper time for a break to avoid participants ’ fatigue. The detailed EEG /fMRI schedule is illustrated in Figure 1.

**Figure 1.**
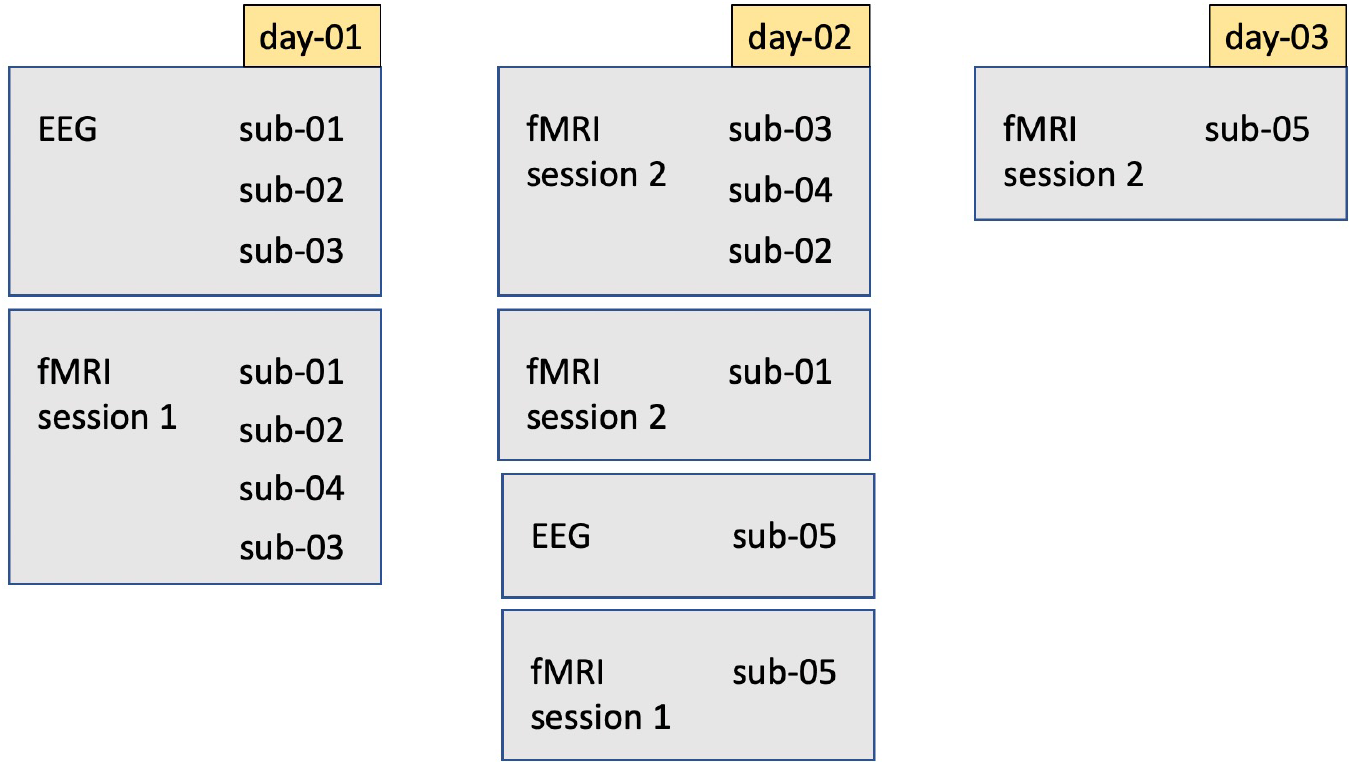
EEG /fMRI schedule - The overall experimental protocol performed over a period of 3 consecutive days. Day-01 contains the EEG recordings as well as the recordings of the first fMRI session of sub-01, sub-02, and sub-03. Note that also the fMRI session for sub-04 took place. Day-02 contains the fMRI recordings for session 2 of sub-01, sub-02, sub-03, and sub-04, the fMRI recordings for session 1 of sub-05 and the EEG recordings of sub-05. Day-03 contains the fMRI recordings for session 2 of sub-05. There was always a relaxation period of at least one hour in between the recordings and proper time for a break to avoid participants ’ fatigue.

The experimental protocol for both modalities (fMRI and EEG) was designed using E-Prime 3.0^46^ and is illustrated in Figure 2. Huth *et al*. ^16^ shows that semantically selective brain areas appear to be organised in the same manner across individuals and provides word frequency statistics for the text corpus employed. Based on the reference study, two categories, social and number, with four words each were selected. The two selected categories were mapped into different brain areas and the selected words appear to have a high word co-occurrence frequency. The social category contained the words *child, daughter, father, and wife*. The number category contained the words *four, three, ten, and six*. The textual representation of the words was presented randomly on the screen in front of the participant. There were a total of 2,080–2,200 fMRI volumes collected per subject, divided into two sessions. Each volume contained 100×100×68 voxels. The EEG recordings provided a total of 320×64×1,024 samples per subject.

**Figure 2.**
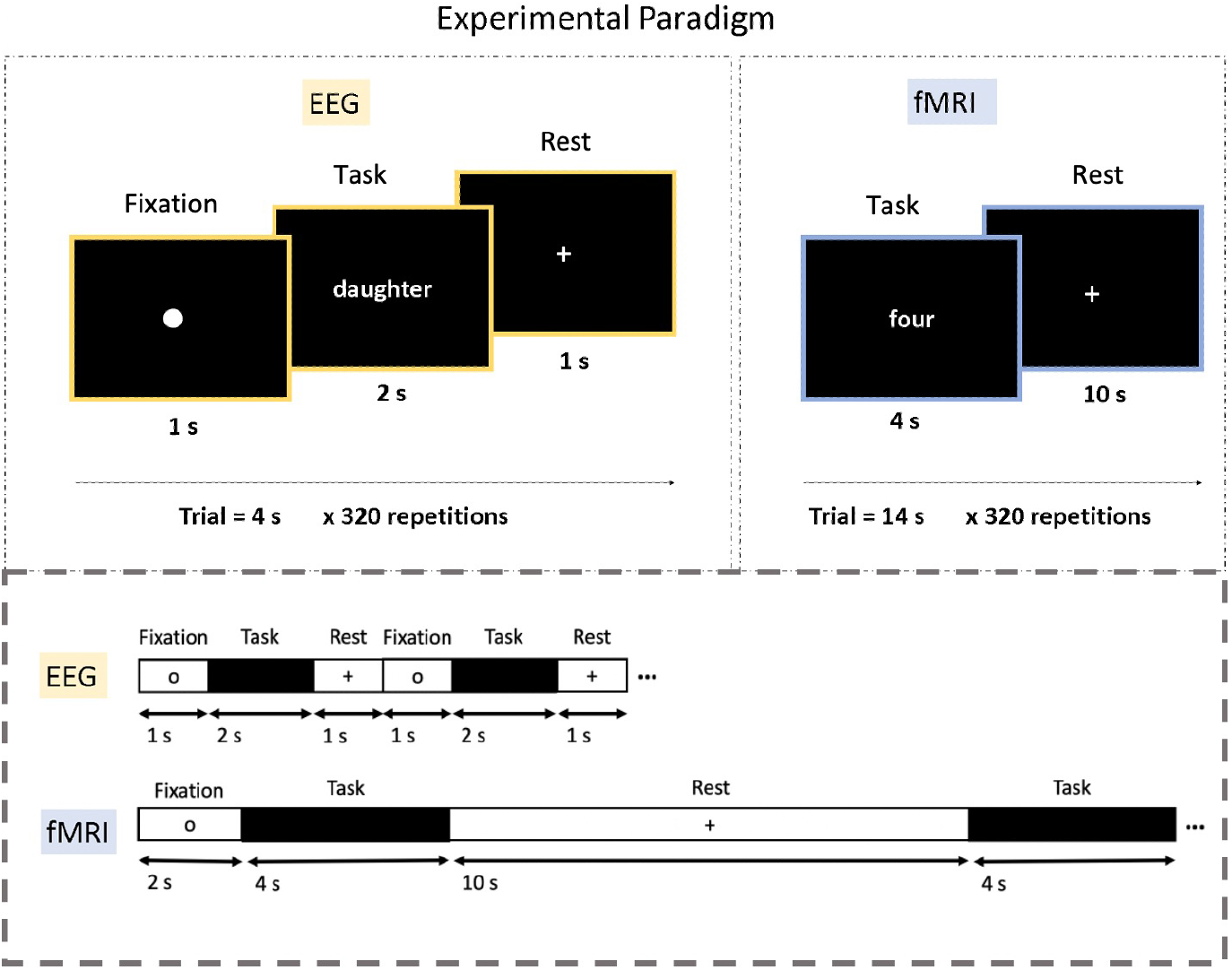
Experimental paradigm - The trial designs are depicted in the top part of the figure and the respected timelines for the trials are shown at the bottom. During the fMRI inner-speech task, participants were requested to think of the presented word as many times as possible. There were 320 trials in total, split into two sessions of 160 trials each. For the EEG recordings, there was only one session, and the presented stimulus was imagined only once for each inner-speech task interval. Note that the rest period for the fMRI protocol was longer than that for the EEG protocol

### fMRI procedure

The fMRI recordings consisted of two sessions performed over a period of three days. At the beginning of each session, written instructions for the experiment were presented on the screen until the participants informed the fMRI operator through an intercom that they are ready to proceed with the experiment. A fixation period of 2 s was followed, in which the participants were instructed to fixate their eyes on the centre of the screen. Then, each trial consisted of the inner-speech task (4 s) and a subsequent rest period (10 s). Eight different words were used for the inner-speech task, divided into 2 categories (social or number words), and there were 20 trials for each word in each session; thus, each session consisted of 160 trials. During the inner-speech task, the word stimulus was presented in white font against a black background for 4 s, and the participants were encouraged to repeat the given word in their minds as many times as possible (approximately 4 times) without any accompanying articulation or muscle movement (i.e., using their inner speech). The word stimuli were presented in a randomized order over the 160 trials. During the rest period, a white fixation cross was presented for 10 s, and the participants were allowed to relax and prepare for the next trial. The total duration of the recordings for the 320 repetitions was 74.6 min per participant.

### EEG *procedure*

The EEG recordings consisted of one session with 40 trials per word, using the same stimuli as in the fMRI protocol, that was performed before the fMRI acquisition for all subjects but for sub-05 due to time constraints of the recording facility. At the beginning of the session, the written instructions for the experiment were presented on a screen to the subject until they pressed the spacebar to start the experiment. Each trial included fixation, task, and rest periods, with durations of 1 s, 2 s, and 1 s, respectively. During the fixation period, the participant was instructed to direct their gaze to the centre of the screen, where a small circular fixation point was located. During the task period, the word stimulus was presented for 2 s, and the participants were asked to repeat the stimulus in their minds without any accompanying articulation or muscle movement (i.e., using their inner speech). During the rest period, the participants were allowed to relax and prepare for the next trial. The total duration of the recording, which contained 320 repetitions, was 21.33 min per participant. Note that the rest period for the fMRI protocol was longer than that for the EEG protocol because the fMRI BOLD signal typically peaks approximately 5 s after stimulus onset and takes approximately 14 s to recover to baseline levels^47^.

### fMRI Preprocessing

The fMRI data were preprocessed with SPM12^48^. First, spatial displacement maps were calculated for each session. These were used for motion correction of the functional data. Slice-timing correction was performed as the fMRI data were acquired in an interleaved order. Next, images were coregistered to the T1-weighted structural scan with a normalized mutual information cost function. Prior to normalization, these images were used for within-subject classification.

To verify that neural activity related to inner speech and the two semantic categories (social and number words) was as expected, further processing was conducted (see “fMRI activation – group level”). The origin was manually set to the anterior commissure, followed by normalization to the Montreal Neurological Institute (MNI) space. Smoothing using an 8-mm full-width at half-maximum (FWHM) Gaussian kernel was applied. We estimated a general linear model (GLM) convolved with a canonical haemodynamic response function. The category regressors (social or number word) were time-locked to the onset of its respective inner-speech word with a duration of 4 s. The rest of the regressors had durations of 10 s. Nuisance variables, such as movement parameters calculated in the previous realignment step, were also included. This GLM enabled investigation of the activation at the subject level; subsequently, a fixed effect analysis was applied to determine activation at the group level. The planned comparisons included inner speech and rest (inner speech – rest) and the stimulus category (number word – social word; social word – number word). For all analyses, the extent cluster threshold was *^K^E* > 20 with a familywise error (FWE) correction of *p* < *0.05* at the voxel level.

### EEG preprocessing

In the current work, we utilized EEGLAB^49^ to preprocess the EEG data. EEGLAB is a MATLAB toolbox for processing continuous and event-related EEG, MEG, and other electrophysiological data. The toolbox has features such as independent component analysis (ICA), time/frequency analysis, artefact rejection, event-related statistics, and several useful visualization modes for averaged and single-trial data.

The raw BioSemi EEG data in .bdf format were imported to EEGLAB using reference channel 48 (Cz). A multitude of internal and environmental causes can generate temporal drifts, which change over time and across the electrodes. To reduce the impact of such variances, it is usual practice to perform a so-called baseline correction. In this study, baseline correction was applied using a zero-phase finite impulse response (FIR) high-pass filter at 0.1 Hz. Low-frequency and high-frequency signals, which are commonly caused by environmental/muscle noise in scalp EEG and are not usually the focus of analysis, were filtered out.

Noise below and above a given frequency was retained using low-pass and high-pass filtering. Here, we applied zero-phase finite impulse response (FIR) bandpass filtering with 0.1 Hz (lower edge) and 50 Hz (higher edge) boundaries of the frequency bandpass, eliminating the requirement for a notch filter. Rereferencing facilitates data cleaning by providing an estimate of physiological noise at baseline. In this study, rereferencing was performed using the average reference to Cz, excluding channels 65-70 (the mastoid and ocular electrodes); averaging referencing was chosen over rereferencing from the mastoid electrodes to guard against introducing any signal artifacts which may have resulted from differences in placement of the external electrodes between participants (as per, for example, our decision to disregard the data from subject 4 (sub-04), to ensure overall data integrity).

Channels were manually inspected, and bad channels were rejected and not interpolated. Time-locked epochs were extracted using start-stop limits fixed within the interval [0, 2 s]. ICA can be used to identify data segments strongly influenced by motor-related artefacts, such as eye blinking and movement of the jaw, neck, arm, or upper back, for removal. Figure 3 illustrates two out of the 64 ICA components. In this study, we examined the topography as well as the spectrogram and frequency variation to decide whether a component should be retained. In examining the topographies, high activity in the far-frontal projections is a strong indicator of electrooculography (EOG) artifacts; in the spectra, decreasing power with a slope that is more shallow and spread more evenly over the frequency range is also a strong indicator of an EOG artifact: taken together (along with the ocular reference channels, 67-70, which serve to provide the EOG signal pattern that the independent components are matched against) we reliably identified artifact-related independent components to zero-out from the data, as part of a manual inspection and cleaning process that is often more reliable than algorithmic methods relying on peak-to-peak signal information, which may vary greatly between subjects.

**Figure 3.**
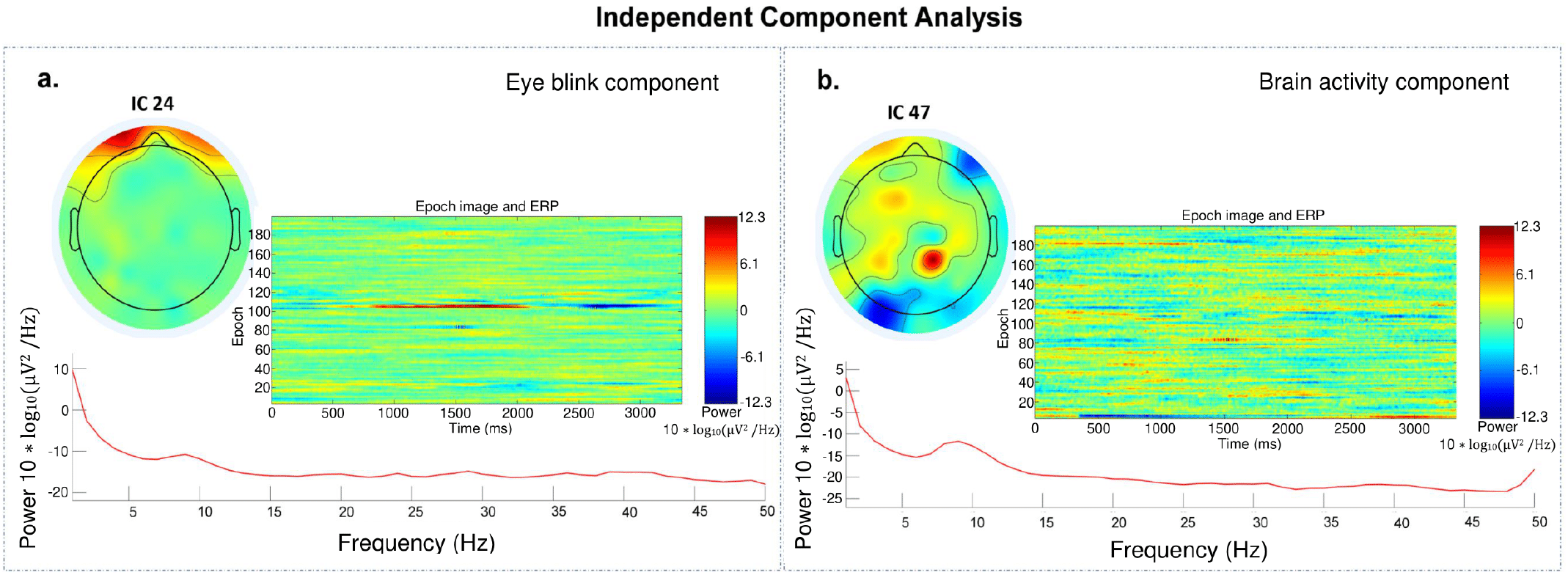
Components decomposed by ICA, showing eye blinks and brain activity. Top left: topography map illustrating the projection of the independent component activity. Center: event-related potentials of the independent component and its associated power, per epoch. Bottom: spectrum of the independent component.

Finally, we extracted epochs according to the *time-locking event*. We performed this process manually, with the epoch limit [0,2 s], this interval encapsulated the main activity.

### Data Records

The anonymized EEG and fMRI data of the four subjects are available in *Brain Imaging Data Structure* (BIDS) format (https://bids-specification.readthedocs.io/en/stable/) at the OpenNeuro repository^50^. The data for each subject are organized into three sessions: two for the fMRI modality (ses-01 and ses-02) and one for the EEG modality (ses-EEG), as shown in Figure 4. A total of 2,560 trials are provided in this dataset, out of which half (1,280) were with fMRI and half were with EEG. There are 4 directories, one for each subject (i.e., sub-01, sub-02, sub-03, and sub-05). The data from sub-04 were deemed unfit for use and thus were not made available.

**Figure 4.**
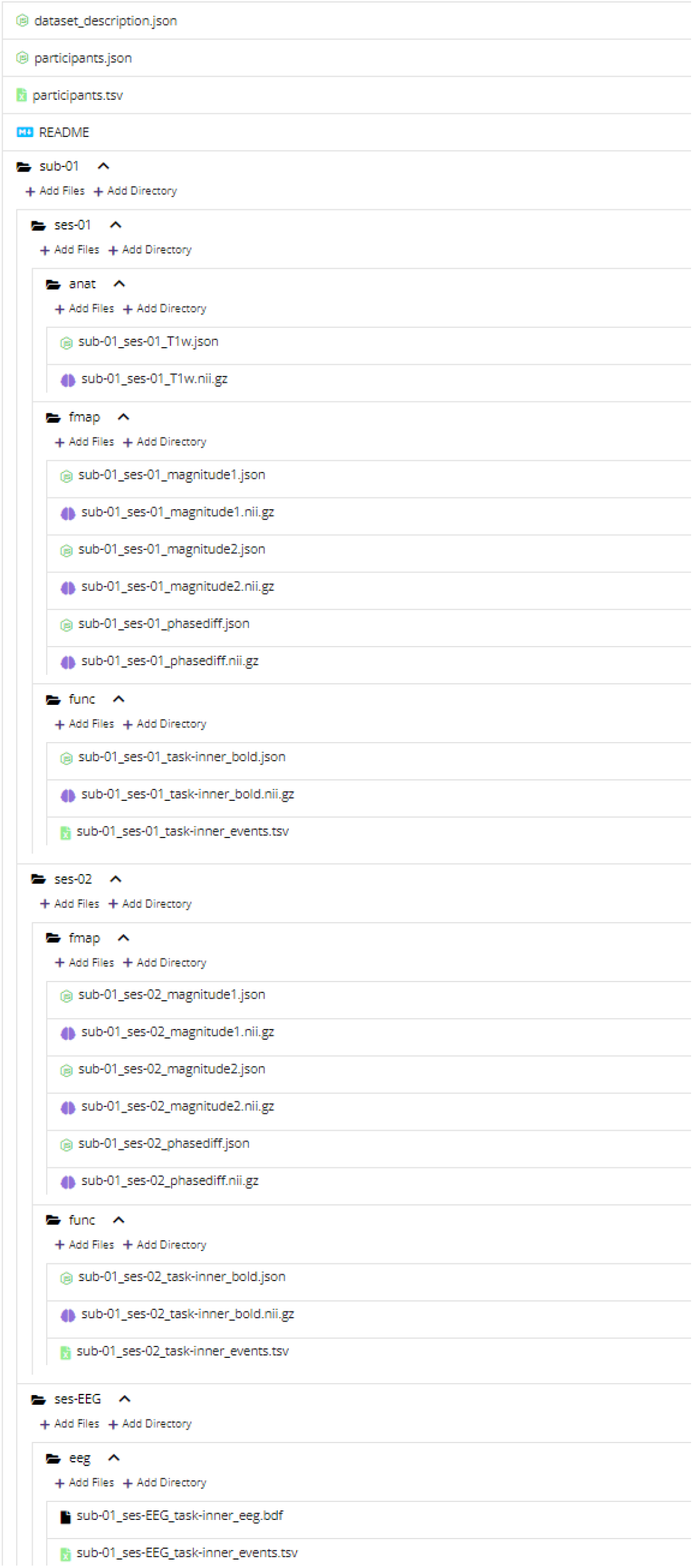
Example dataset structure for subject 1 (sub-01). The data are organized into three sessions: two for the fMRI modality (ses-01 and ses-02) and one for the EEG modality (ses-EEG).

#### fMRI

All Digital Imaging and Communications in Medicine (DICOM) files with fMRI data were converted into Neuroimaging Informatics Technology Initiative (NIFTI) format using *MRIcroGL v1.2.0211006* (https://www.nitrc.org/plugins/mwiki/index.php/mricrogl) and then organized into BIDS format. Regarding file organization, each individual subject folder contains the three sessions described above. The ses-01 folder contains three subfolders that include the anatomical (anat), field map (fmap), and functional (func) images. The ses-02 folder contains two subfolders, namely, fmap and func, as anatomical scans were not performed in session 2 for any of the subjects. Each .nifti file in the dataset is accompanied by the corresponding .json file. The anatomical image from session 1 of each subject consists of the anatomical scan with file names in the following format: sub-*XX*-*YY*-T1W, where *XX* denotes the subject ID and *YY* denotes the session ID. The fmap folder in each session of a subject consists of three .nifti files and their corresponding .json files. Two of the .nifti files are for magnitude, and one is for phase difference. The two magnitude files are named with the following format: sub-*XX*-*YY*-magnitudeZ, where *XX* denotes the subject ID, *YY* denotes the session ID, and Z takes the value 1 for *TE*1 = 4.92ms and 2 for *TE*2 = 7.38ms. The .nifti file for the phase difference image is named with the following format: sub-*XX*-ses-*YY*-phasediff, where *XX* denotes the subject ID and *YY* denotes the session ID. The functional data are made available in the func folder, where the .nifti file and its corresponding .json file have the following format: sub-*XX*-ses-*YY*-task-inner-bold, where *XX* denotes the subject ID and *YY* denotes the session ID. The task event file is also available as a .tsv file named sub-XX-ses-YY-task-inner-events, where *XX* denotes the subject ID and *YY* denotes the session ID in the corresponding func folders.

#### EEG

The EEG data were collected in one session, and raw EEG files are available in the ses-EEG folder for each subject. The raw EEG data are available in .bdf format. The .bdf file was exported using EEGLAB software v2021.1, and the sampling rate was 512. Each event and its corresponding ID and description are presented in Table 2. The different channel data are provided in 72 rows (64 EEG, 8 external). Out of 8 external electrodes, only six were connected; specifically, the CMS and DRL electrodes were not recorded.

**Table 2.** Event IDs and their accompanying descriptions.

| EventID | Description |
| --- | --- |
| 1 | fixation |
| 2 | rest |
| 111 | child |
| 112 | daughter |
| 113 | father |
| 114 | wife |
| 125 | four |
| 126 | three |
| 127 | ten |
| 128 | six |

The task events are provided in a .tsv file for corresponding subjects in the respective ses-EEG folders. The continuous recordings of the 64 EEG channels, the 6 external channels, and the labeled events were included in the saved files.

### Technical Validation

#### EEG

An event related potential (ERP) is the measured brain activity in response to a stimulus, therefore suitable for verifying the technical validity of the data. Although the entire dataset is available at the OpenNeuro repository^50^, in this work, we provide the ERP activity for two subjects in Figure 5, as an example. In order to represent all available genders in this study, sub-01 (male) and sub-03 (female) were selected.

**Figure 5.**
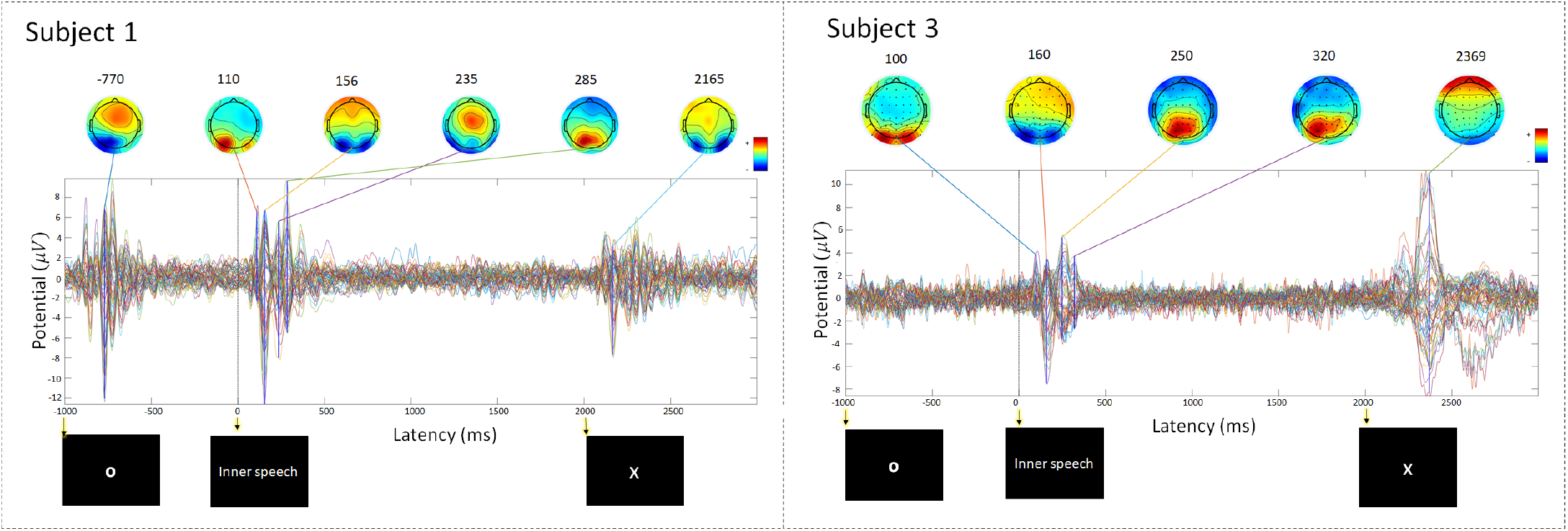
Plot of event-related potentials and topographical maps of activity for subjects 1 (sub-01) and 3 (sub-03). Both plots are created from preprocessed data by averaging over 320 trials for each subject. The 64 coloured waves correspond to the 64 EEG channels. The time axis shows the duration of a single trial and corresponds to a total duration of 4,000 ms. Notably, the time axis starts from a negative value of t=-1,000 ms, which corresponds to the 1,000 ms fixation period at the beginning of a trial. The times marked with arrows indicate the start of the fixation, inner-speech task, and rest periods (at −1,000 ms, 0 ms and 2,000 ms respectively.

From both plots, it can be observed that more pronounced potential deflections occur after stimulus presentation (i.e., after t=0 ms); however, a small range of deflection can also be noticed within the two other periods (rest and fixation); the presentation of new visual stimuli to the subjects, to signal the start of each period within each trail, has resulted in these evoked responses (so-called fixation-onset ERPs^51^).

The activity at 235 ms and 285 ms for sub-01, and at 250 ms and 320 ms for sub-03 indicate prominent brain activity stimulus onset. Specifically, sub-01 displayed high activity, linked to the frontal lobe, during the first 500 ms compared to the low activity of sub-03 in the same period. This difference might be a result of eye movements by sub-01. Interestingly, for sub-03, the plotted waves were close to the baseline during the fixation period, indicating low brain activity associated with gaze at the fixation circle. Data from both subjects followed similar pattern during the inner-speech task (0-500 ms). As shown in Figure 5, strong positive and negative deflections occurred during 0-500 ms, which indicate the evoked response upon presentation of the visual stimulus, and increased brain activity during the performance of the inner-speech task. These evoked responses indicate strong P300 and N400 components, which are observed in similar trials for imagined speech^52^ and validate the inner-speech data that have been recorded under our experimental protocol.

Next, to provide a more detailed analysis of the activated brain areas, we generated topological maps (after ICA decomposition). Figure 6 depicts the activation in response to stimuli in the two semantic categories (number or social words) for all subjects in topological maps. As shown in the figure, it appears that these activities were mainly dominated by frontal and central regions. Notably, the activity in response to all stimuli (eight words) is shown in the topological maps in Figure 7. In this figure, 4 ICs with high brain activity are shown for all words. For a specific IC, regions were differently activated for each of the eight words. The topographic projections of each word illustrate the average difference in brain activity between the inner-speech task of each word; the variance between these projections confirms the subject’s different activation regions, both between and within our semantic categories, which further validates the data.

**Figure 6.**
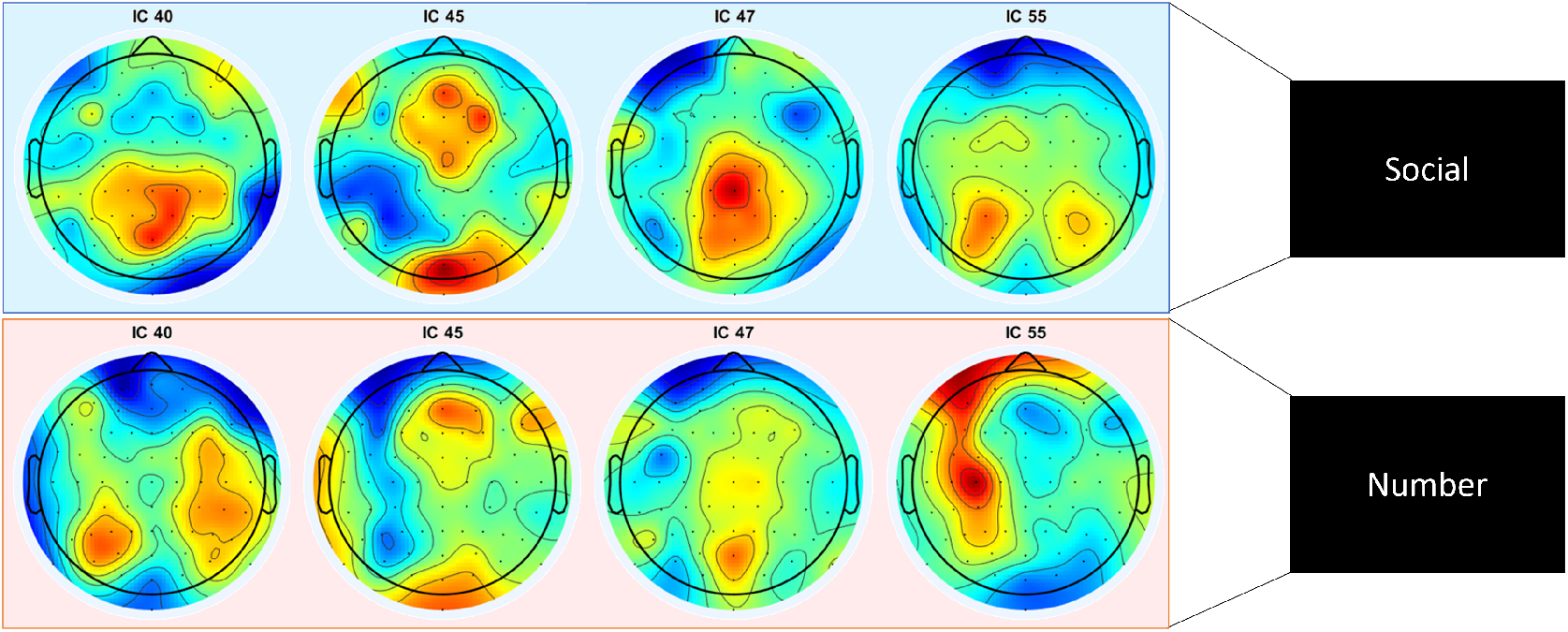
Topological maps (averaged over all subjects) corresponding to stimuli from the number and social categories for components 40, 45, 47, and 55.

**Figure 7.**
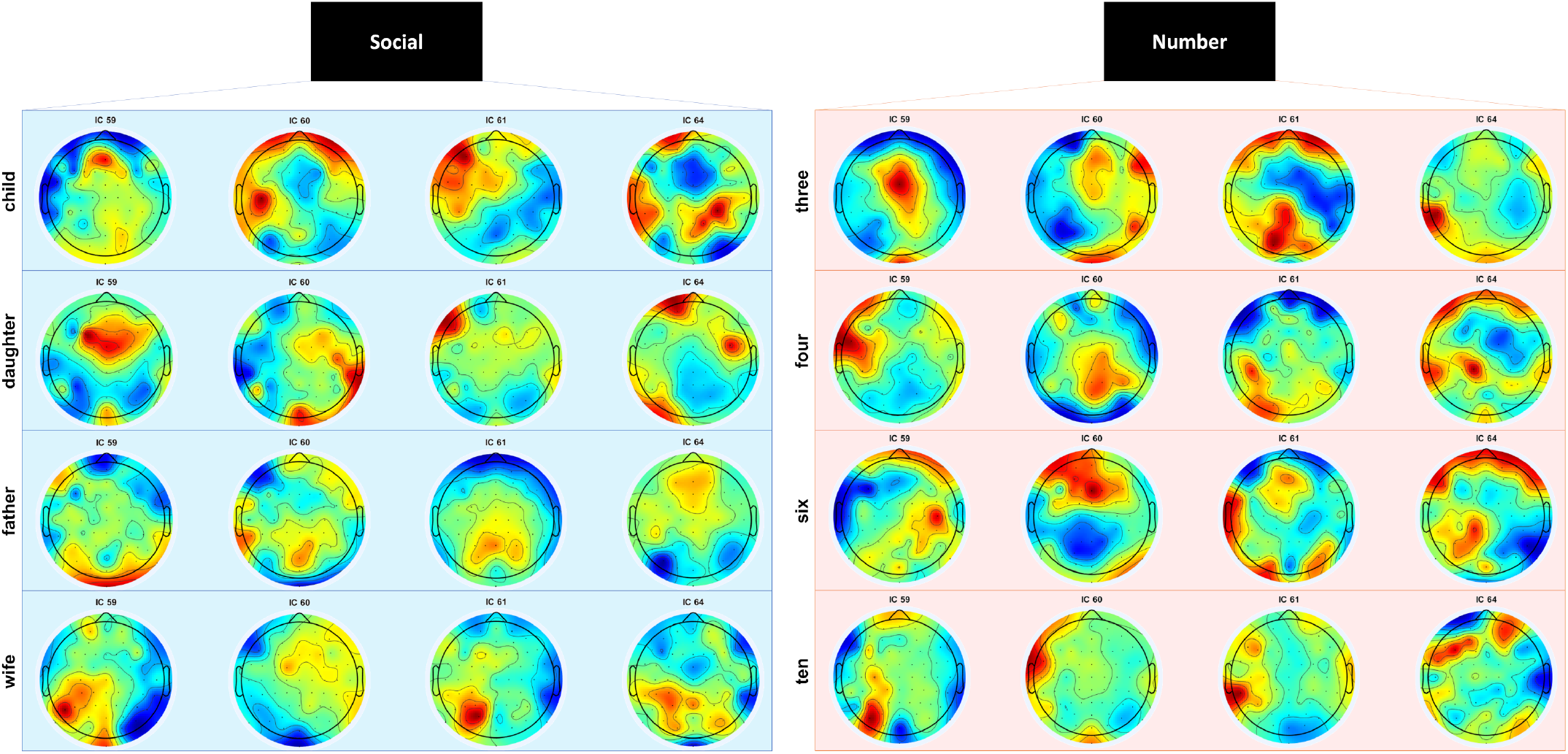
Topological maps (averaged over all subjects) for each stimulus in the number and social categories for components 59, 60, 61, and 64.

### fMRI activation - group level

In this study, the group level analyses were conducted to verify that the inner-speech task activated neural regions connected to inner speech. As expected, the inner-speech task activated language- and orthographic-related regions when compared with the baseline rest condition. The increased activity during inner speech (see Figure 8**(a)** and Table 3) is indicated by average BOLD activation displaying significant activation in areas directly related to language processing, including Wernicke’s area (Brodmann’s area (BA) 22) in the left hemisphere and Broca’s area and surrounding regions. Further activation was found in the left supramarginal gyrus, which (alongside the pars opercularis) has been implicated in inner speech^53^, and the angular gyrus, which is related to semantic processing^54, 55^. Areas of high activation also included visual processing regions, such as the bilateral secondary visual cortex and the right frontal eye fields, and orbitofrontal area. These regions likely relate to processing of visual word forms, as the word cue was presented orthographically. Not surprisingly, motor regions, including the primary motor cortex and premotor and supplementary motor cortices, were activated by the inner-speech task. Motor activity still occurs during inner-speech^56^, albeit at reduced activation levels compared to outer speech (spoken aloud)^57^. These findings support the reliability of the data as they indicate that the inner-speech task activated language, orthographic, and motor related regions.

**Figure 8.**
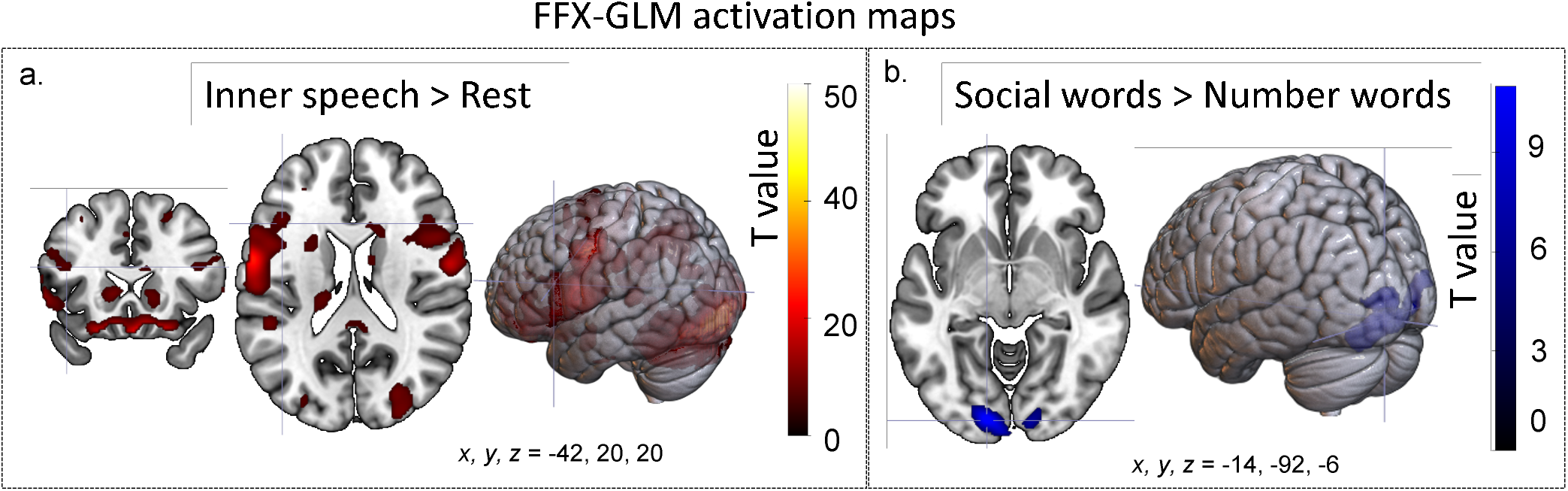
Activation map generated from fixed effects (FFX) and the general linear model (GLM) for the following: a. areas more highly activated during the inner-speech task than the resting condition (number and social words combined) – this slice is selected to highlight activation in Broca’s area (coordinates: −42, 20, 20), and b. areas more highly activated by social words than number words – this slice is selected to highlight activation in the secondary visual cortex (coordinates: 14, −92, −6). An increase in activation was found for the reverse contrast (i.e., increased activation for number words over social words).

**Table 3.**
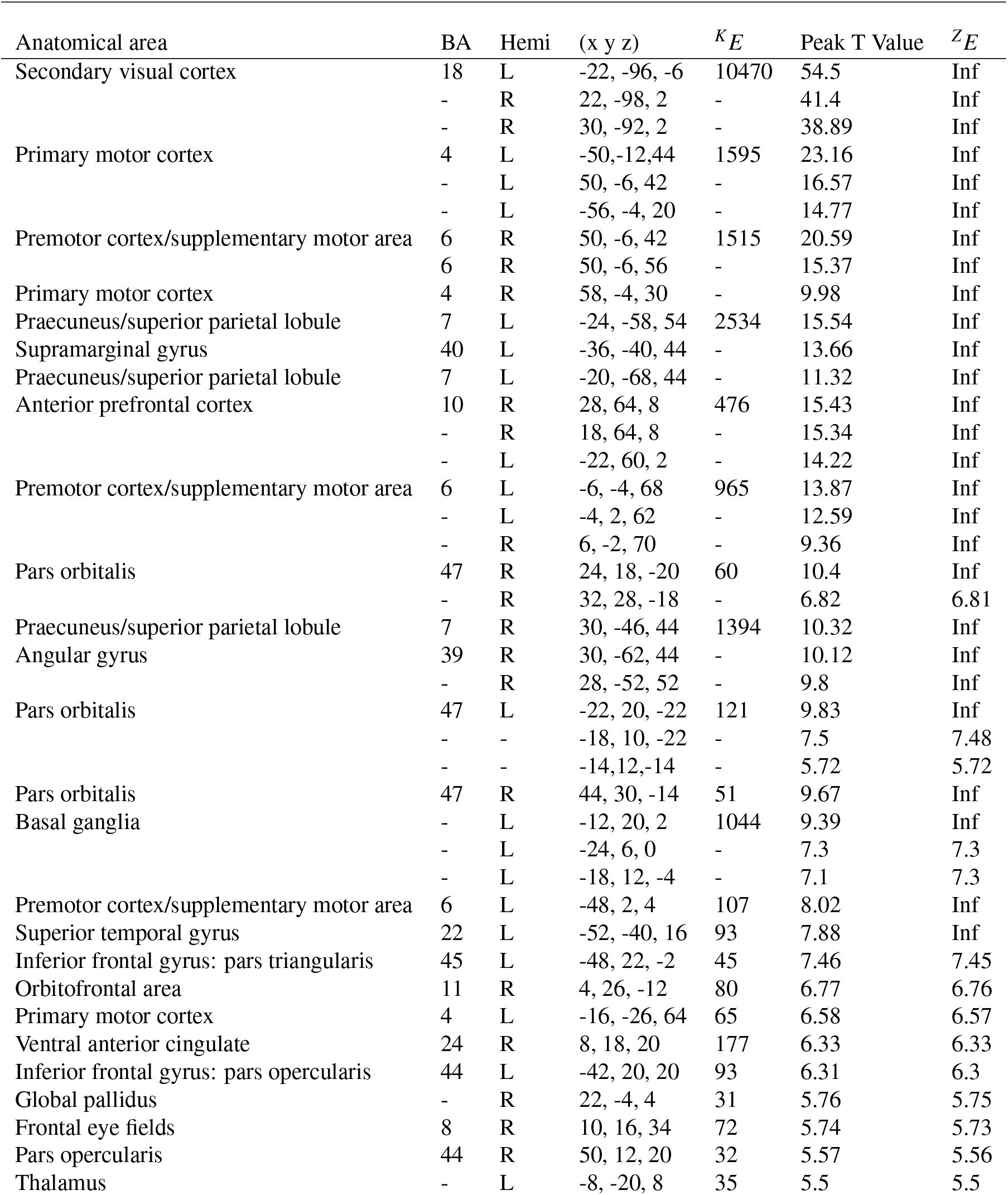
Areas with increased activation during inner speech relative to baseline (the rest condition). Coordinates are in Montreal Neurological Institute (MNI) space. *BA* = Brodmann’s Area, *Hemi* = hemisphere, *^K^E*= cluster size. The cluster threshold was set to 20, with family-wise error (FWE)-adjusted *p*<0.05

Comparison of the two semantic categories (social vs. number words) revealed that social words elicited more activation than number words in the bilateral secondary visual cortex and the right primary visual cortex (see Table 4 and Figure 8(b)). No areas showed significantly higher activation for number words compared to social words.fMRI activation - individual subjects

**Table 4.**
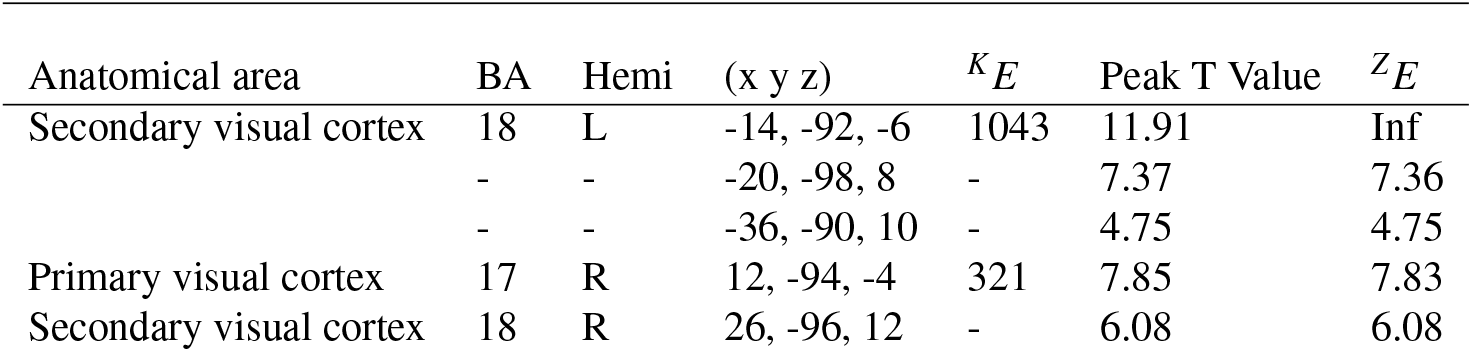
Areas with increased activation for social words relative to number words during inner speech, i.e. the baseline. Coordinates are in Montreal Neurological Institute (MNI) space. *BA* = Brodmann’s area, *Hemi* = hemisphere, *^K^E* = cluster size. The cluster threshold was set to 20. The family-wise error (FWE) was adjusted to *p*<0.05

Decoding studies are often conducted within rather than between subjects due to the large extent of individual variance in neural anatomy and functional activity. Therefore, we provided subject-level results to enable researchers to select among subjects based on brain activation profiles. As seen in Figure 9, the results are consistent across subjects.

**Figure 9.**
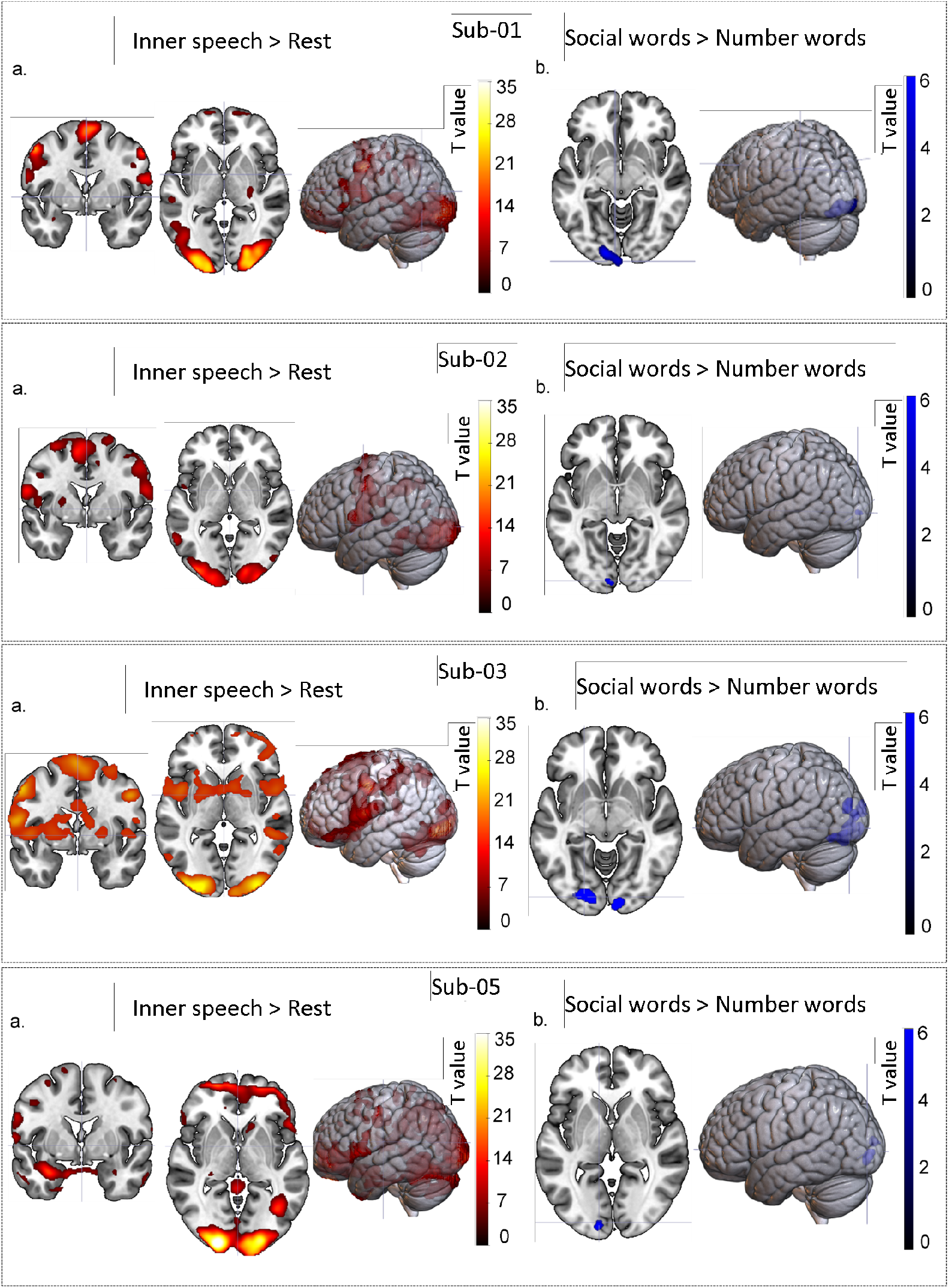
BOLD signals of each subject. **a.** Area more highly activated during the inner-speech task in the rest condition (numbers and social words combined). **b.** Areas more highly activated by social words than number words (no areas were more highly activated by number words than social words).

### fMRI framewise displacement

Motion-related artifacts can compromise data quality. Frames that are contaminated with motion above a certain threshold can be rejected by calculating head motion artifacts through framewise displacement (FD). FD is an overall estimate of movement over time for each subject, which incorporates subtle in-scanner movements. We calculated the FD for each subject and session in Nipype, according to Power *et al*.^58^. The average FD (in mm) across frames for each subject was as follows: sub-01, session 1=0.13 and session 2=0.14; sub-02, session 1=0.15 and session 2=0.14; sub-03, session 1=0.11 and session 2=0.1; and sub-05, session 1=0.2 and session 2=0.22; see Figure 10. There was rarely motion exceeding the size of one voxel (2 × 2 × 2 mm). The analysis also showed that all subjects had a mean FD under 0.25 mm; however, researchers may choose to omit specific volumes with FD values higher than 0.5 mm.

**Figure 10.**
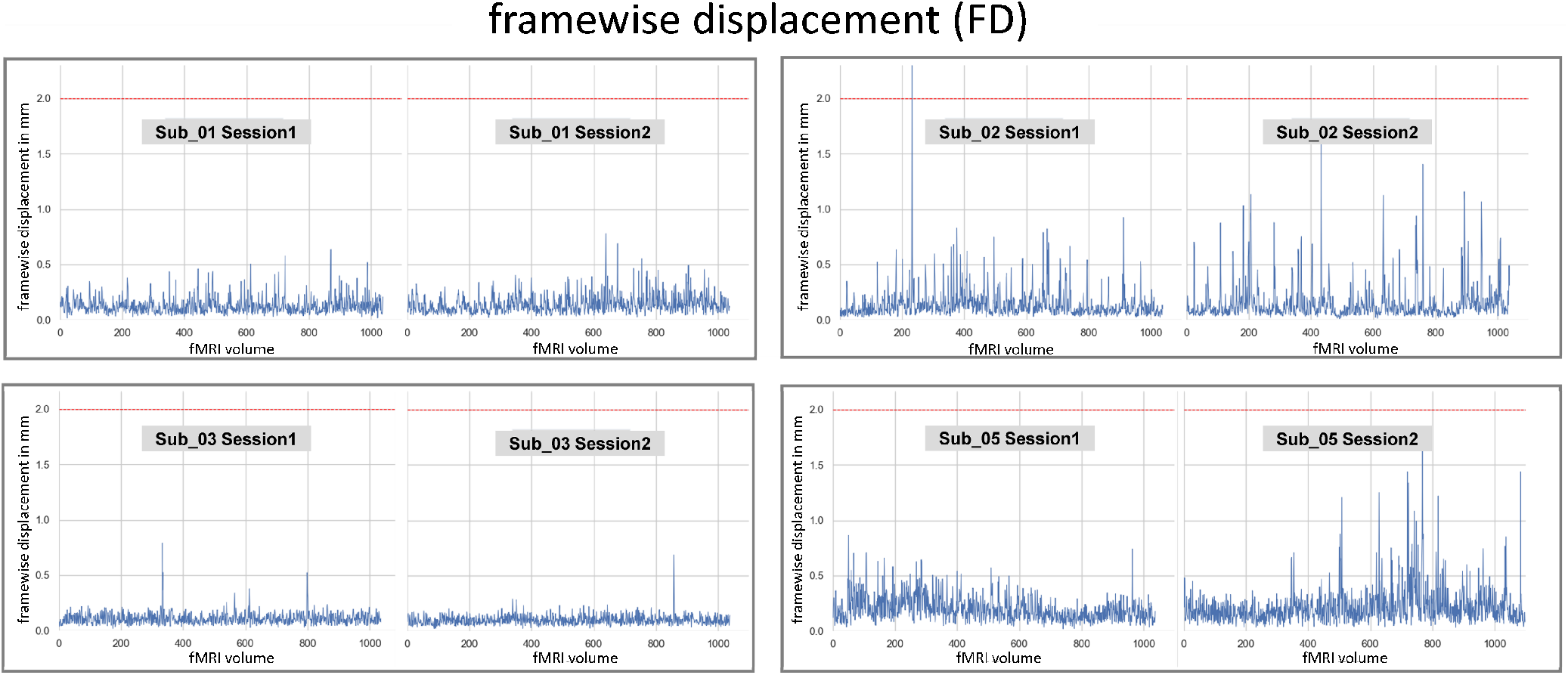
The framewise displacement (FD; in mm) calculated across each subject and session is shown across each volume. The red dashed line indicates the voxel size of the functional images (2×2×2 mm).

## Code Availability

The code used to preprocess the EEG and fMRI data as well as the two stimulation protocols (one for each modality) are publicly available at: https://github.com/LTU-Machine-Learning/Inner_Speech_EEG_FMRI.

## Acknowledgements

This research was funded by the Grants for Excellent Research Projects Proposals of SRT.ai 2022. We would like to thank the Stockholm University Brain Imaging Centre (SUBIC) and, in particular, Rita Almeida and Patrik Andersson for giving us access to their facilities and for supporting us in this endeavour. We thank Petter Kallioinen and Christoffer Schiehe-Forbes for their valuable support during the acquisition of EEG data. Finally, we would also like to thank all participants for taking part in this study.

## Author contributions statement

F.S.L.: project conception, experimental design, data acquisition, writing, and finalizing the manuscript. V.G.: conducted the processing part of the EEG data, and writing EEG-related sections of the manuscript. R.S.: processing of EEG data and writing EEG-related sections of the manuscript. K.D.: processing and technical validation of fMRI data and writing all sections of the manuscript. N.A.: review of all sections of the manuscript. S.R.: data management and availability. S.W.: advised the pipeline for EEG data processing and validation, processing results, and writing related sections of the manuscript. H.W.: designed the pipeline for processing and technical validation of fMRI data as well as writing all sections of the manuscript. M.L.: advised the experimental and research design and the writing of the manuscript. J.E.: advised the experimental design, technical validation of fMRI data, and writing of the manuscript. All authors reviewed the manuscript.

## References

1. He, B., Yuan, H., Meng, J. & Gao, S. Brain–computer interfaces. In Neural engineering, 131–183 (Springer, 2020).

2. Abiri, R., Borhani, S., Sellers, E. W., Jiang, Y. & Zhao, X. A comprehensive review of eeg-based brain–computer interface paradigms. J. neural engineering 16, 011001 (2019).

3. Alderson-Day, B. & Fernyhough, C. Inner speech: development, cognitive functions, phenomenology, and neurobiology. Psychol. bulletin 141, 931 (2015).

4. Whitford, T. J. et al. Neurophysiological evidence of efference copies to inner speech. Elife 6, e28197 (2017).

5. Smallwood, J. & Schooler, J. W. The science of mind wandering: empirically navigating the stream of consciousness. Annu. review psychology 66, 487–518 (2015).

6. Filik, R. & Barber, E. Inner speech during silent reading reflects the reader’s regional accent. PloS one 6, e25782 (2011).

7. Angrick, M. et al. Real-time synthesis of imagined speech processes from minimally invasive recordings of neural activity. Commun. biology 4, 1–10 (2021).

8. Dash, D., Ferrari, P., Berstis, K. & Wang, J. Imagined, intended, and spoken speech envelope synthesis from neuromagnetic signals. In International Conference on Speech and Computer, 134–145 (Springer, 2021).

9. McGuire, P. et al. The neural correlates of inner speech and auditory verbal imagery in schizophrenia: relationship to auditory verbal hallucinations. The Br. J. Psychiatry 169, 148–159 (1996).

10. Barber, L., Reniers, R. & Upthegrove, R. A review of functional and structural neuroimaging studies to investigate the inner speech model of auditory verbal hallucinations in schizophrenia. Transl. psychiatry 11, 1–12 (2021).

11. Perrone-Bertolotti, M., Rapin, L., Lachaux, J.-P., Baciu, M. & Loevenbruck, H. What is that little voice inside my head? inner speech phenomenology, its role in cognitive performance, and its relation to self-monitoring. Behav. brain research 261, 220–239 (2014).

12. Skeide, M. A. & Friederici, A. D. The ontogeny of the cortical language network. Nat. Rev. Neurosci. 17, 323–332 (2016).

13. Blank, S. C., Scott, S. K., Murphy, K., Warburton, E. & Wise, R. J. Speech production: Wernicke, broca and beyond. Brain 125, 1829–1838 (2002).

14. Sahin, N. T. et al. Sequential processing of lexical, grammatical, and phonological information within broca’s area. Science 326, 445–449 (2009).

15. Mitchell, T. M. et al. Predicting human brain activity associated with the meanings of nouns. science 320, 1191–1195 (2008).

16. Huth, A. G. et al. Natural speech reveals the semantic maps that tile human cerebral cortex. Nature 532, 453–458 (2016).

17. Rueckl, J. G. et al. Universal brain signature of proficient reading: Evidence from four contrasting languages. Proc. Natl. Acad. Sci. 112, 15510–15515 (2015).

18. Wilson, J. A., Felton, E. A., Garell, P. C., Schalk, G. & Williams, J. C. Ecog factors underlying multimodal control of a brain-computer interface. IEEE transactions on neural systems rehabilitation engineering 14, 246–250 (2006).

19. Fabiani, G. E., McFarland, D. J., Wolpaw, J. R. & Pfurtscheller, G. Conversion of eeg activity into cursor movement by a brain-computer interface (bci). IEEE transactions on neural systems rehabilitation engineering 12, 331–338 (2004).

20. Andersson, P. et al. Real-time decoding of brain responses to visuospatial attention using 7t fmri. PloS one 6, e27638 (2011).

21. Kamavuako, E. N., Sheikh, U. A., Gilani, S. O., Jamil, M. & Niazi, I. K. Classification of overt and covert speech for near-infrared spectroscopy-based brain computer interface. Sensors 18, 2989 (2018).

22. Rezazadeh Sereshkeh, A. et al. Development of a ternary hybrid fNIRS-EEG brain–computer interface based on imagined speech. Brain-Computer Interfaces 6, 128–140 (2019).

23. Dash, D. et al. Meg sensor selection for neural speech decoding. IEEE Access 8, 182320–182337 (2020).

24. Dash, D. et al. Decoding imagined and spoken phrases from non-invasive neural (MEG) signals. Front. neuroscience 14 (2020).

25. Aggarwal, S. & Chugh, N. Signal processing techniques for motor imagery brain computer interface: A review. Array 1, 100003 (2019).

26. Chholak, P. et al. Visual and kinesthetic modes affect motor imagery classification in untrained subjects. Sci. reports 9, 1–12 (2019).

27. Donchin, E., Spencer, K. M. & Wijesinghe, R. The mental prosthesis: assessing the speed of a p300-based brain-computer interface. IEEE transactions on rehabilitation engineering 8, 174–179 (2000).

28. da Silva-Sauer, L., Valero-Aguayo, L., de la Torre-Luque, A., Ron-Angevin, R. & Varona-Moya, S. Concentration on performance with p300-based bci systems: A matter of interface features. Appl. ergonomics 52, 325–332 (2016).

29. Herff, C. et al. Brain-to-text: decoding spoken phrases from phone representations in the brain. Front. neuroscience 9, 217 (2015).

30. Martin, S., Iturrate, I., Millán, J. d. R., Knight, R. T. & Pasley, B. N. Decoding inner speech using electrocorticography: Progress and challenges toward a speech prosthesis. Front. neuroscience 12, 422 (2018).

31. Panachakel, J. T. & Ramakrishnan, A. G. Decoding covert speech from eeg-a comprehensive review. Front. Neurosci. 392 (2021).

32. Cooney, C., Folli, R. & Coyle, D. Optimizing layers improves CNN generalization and transfer learning for imagined speech decoding from EEG. In 2019 IEEE International Conference on Systems, Man and Cybernetics (SMC), 1311–1316 (IEEE, 2019).

33. Schirrmeister, R. T. et al. Deep learning with convolutional neural networks for EEG decoding and visualization. Hum. brain mapping 38, 5391–5420 (2017).

34. van den Berg, B., van Donkelaar, S. & Alimardani, M. Inner speech classification using eeg signals: A deep learning approach. In 2021 IEEE 2nd International Conference on Human-Machine Systems (ICHMS), 1–4 (IEEE, 2021).

35. Yoo, S.-S. et al. Brain–computer interface using fmri: spatial navigation by thoughts. Neuroreport 15, 1591–1595 (2004).

36. Zhao, S. & Rudzicz, F. Classifying phonological categories in imagined and articulated speech. In 2015 IEEE International Conference on Acoustics, Speech and Signal Processing (ICASSP), 992–996 (IEEE, 2015).

37. Coretto, G. A. P., Gareis, I. E. & Rufiner, H. L. Open access database of EEG signals recorded during imagined speech. In 12th International Symposium on Medical Information Processing and Analysis, vol. 10160, 1016002 (2017).

38. Nguyen, C. H. et al. Inferring imagined speech using EEG signals: a new approach using Riemannian manifold features. J. neural engineering 15, 016002 (2017).

39. Ferreira, C. et al. Inner speech in portuguese: Acquisition methods, database and first results. In International Conference on Computational Processing of the Portuguese Language, 438–447 (Springer, 2018).

40. Nieto, N., Peterson, V., Rufiner, H. L., Kamienkowski, J. E. & Spies, R. Thinking out loud, an open-access eeg-based bci dataset for inner speech recognition. Sci. Data 9, 1–17 (2022).

41. Perronnet, L. et al. Unimodal versus bimodal eeg-fmri neurofeedback of a motor imagery task. Front. Hum. Neurosci. 11, 193 (2017).

42. Cooney, C., Folli, R. & Coyle, D. A bimodal deep learning architecture for eeg-fnirs decoding of overt and imagined speech. IEEE Transactions on Biomed. Eng. (2021).

43. Lioi, G. et al. Simultaneous eeg-fmri during a neurofeedback task, a brain imaging dataset for multimodal data integration. Sci. data 7, 1–15 (2020).

44. Berezutskaya, J. et al. Open multimodal ieeg-fmri dataset from naturalistic stimulation with a short audiovisual film. Sci. Data 9, 1–13 (2022).

45. Scrivener, C. L. When is simultaneous recording necessary? a guide for researchers considering combined eeg-fmri. Front. Neurosci. 15, 774 (2021).

46. Schneider, W., Eschman, A. & Zuccolotto, A. E-prime (version 2.0). Comput. software manual]. Pittsburgh, PA: Psychol. Softw. Tools Inc (2002).

47. Dale, A. M. & Buckner, R. L. Selective averaging of rapidly presented individual trials using fmri. Hum. brain mapping 5, 329–340 (1997).

48. Penny, W. D., Friston, K. J., Ashburner, J. T., Kiebel, S. J. & Nichols, T. E. Statistical parametric mapping: the analysis of functional brain images (Elsevier, 2011).

49. Delorme, A. & Makeig, S. EEGLAB: an open source toolbox for analysis of single-trial EEG dynamics including independent component analysis. J. neuroscience methods 134, 9–21 (2004).

50. Liwicki, F. et al. “Bimodal dataset on Inner speech”. OpenNeuro https://doi:10.18112/openneuro.ds004197.v1.0.2 (2022).

51. Katz, C. N. et al. Differential generation of saccade, fixation, and image-onset event-related potentials in the human mesial temporal lobe. Cereb. Cortex 30, 5502–5516 (2020).

52. Villena-González, M., López, V. & Rodríguez, E. Orienting attention to visual or verbal/auditory imagery differentially impairs the processing of visual stimuli. Neuroimage 132, 71–78 (2016).

53. Geva, S. et al. The neural correlates of inner speech defined by voxel-based lesion–symptom mapping. Brain 134, 3071–3082 (2011).

54. Devlin, J. T., Matthews, P. M. & Rushworth, M. F. Semantic processing in the left inferior prefrontal cortex: a combined functional magnetic resonance imaging and transcranial magnetic stimulation study. J. cognitive neuroscience 15, 71–84 (2003).

55. Hartwigsen, G. et al. Dissociating parieto-frontal networks for phonological and semantic word decisions: a condition-and-perturb tms study. Cereb. cortex 26, 2590–2601 (2016).

56. Loevenbruck, H. et al. Neural correlates of inner speaking, imitating and hearing: an fmri study. In ICPhS 2019-19th International Congress of Phonetic Sciences (2019).

57. Palmer, E. D. et al. An event-related fmri study of overt and covert word stem completion. Neuroimage 14, 182–193 (2001).

58. Power, J. D., Barnes, K. A., Snyder, A. Z., Schlaggar, B. L. & Petersen, S. E. Spurious but systematic correlations in functional connectivity mri networks arise from subject motion. Neuroimage 59, 2142–2154 (2012).

